# Rapid cerebrovascular reactivity mapping: Enabling vascular reactivity information to be routinely acquired

**DOI:** 10.1101/094359

**Authors:** Nicholas P. Blockley, James W. Harkin, Daniel P. Bulte

**Author notes:** Address for correspondence: Nicholas Blockley, FMRIB Centre, Nuffield Department of Clinical Neurosciences, University of Oxford, John Radcliffe Hospital, Headington, Oxford, OX3 9DU, UK., Twitter: @nicblockley.

## Abstract

Cerebrovascular reactivity mapping (CVR), using magnetic resonance imaging (MRI) and carbon dioxide as a stimulus, provides useful information on how cerebral blood vessels react under stress. This information has proven to be useful in the study of vascular disorders, dementia and healthy ageing. However, clinical adoption of this form of CVR mapping has been hindered by relatively long scan durations of 7 to 12 minutes. By replacing the conventional block presentation of carbon dioxide enriched air with a sinusoidally modulated stimulus, the aim of this study was to investigate whether more clinically acceptable scan durations are possible. Firstly, the conventional block protocol was compared with a sinusoidal protocol of the same duration of 7 minutes. Estimates of the magnitude of the CVR signal (CVR magnitude) and the relative timing of the CVR response (CVR phase) were found to be in good agreement between the stimulus protocols. Secondly, data from the sinusoidal protocol was reanalysed using decreasing amounts of data in the range 1 to 6 minutes. The CVR magnitude was found to tolerate this reduction in scan duration better than CVR phase. However, these analyses indicate that scan durations in the range of 3 to 5 minutes produce robust data.

## Introduction

Whilst there are many clinical techniques to investigate cerebral physiology at rest, methods to examine the brain during dynamic changes in demand are few in number. Resting perfusion measurement techniques using Computed Tomography (CT) or Magnetic Resonance Imaging (MRI) are able to confirm that the tissue is well supplied at rest, but they do not provide information on how the vasculature will react during increases in demand. By introducing a physiological challenge this aspect of cerebrovascular health can be revealed. There are two main vasoactive challenges that can be used to provide a stress test of the cerebral vasculature: acetazolamide and carbon dioxide. Acetazolamide provides a large physiological challenge by maximally dilating cerebral arterioles. However, its effects are relatively long lasting (∼30 mins) and it therefore has the potential to interfere with the accuracy of other diagnostic measurements made in the same session. In contrast, carbon dioxide may have a weaker vasodilatory effect, but it is rapidly cleared by the lungs enabling multiple short repeated challenges to be performed without interfering with subsequent measurements. When combined with an imaging modality this carbon dioxide stimulus, often termed a hypercapnia challenge, enables cerebrovascular reactivity (CVR) to be mapped at the tissue level. The most common imaging modality for this purpose is MRI. Measurements have been performed using both Arterial Spin Labelling (ASL)^1^ and Blood Oxygenation Level Dependent (BOLD)^2^ weighted techniques, each with their own advantages and disadvantages. BOLD weighted imaging offers higher spatial and temporal resolution and a higher signal to noise ratio (SNR) than ASL, whereas ASL provides quantitative measurements of perfusion change and BOLD does not. In practice BOLD weighted measurements are more commonly used due to their higher sensitivity, more widespread availability and ease of analysis.

The information provided by CVR measurements has proven useful in a range of applications. CVR mapping has perhaps made the biggest impact in the study of vascular disorders of the brain. In patients with unilateral carotid stenosis it has been used to show that CVR is reduced in both the affected and unaffected hemisphere^3^. In Moyamoya patients, who share similar pathological narrowing of cerebral vessels, BOLD based CVR has been shown to give comparable results with ASL based CVR^1^. CVR measurements have also been acquired in difficult patient populations such as in subarachnoid haemorrhage^4^ or in rare conditions such as MELAS^5^ where in both cases only limited biomarkers are available. CVR has also shown promise as a biomarker in dementia and healthy ageing research. In Alzheimer’s disease CVR has been shown to be altered in patients with different variants of the APOE gene^6^. CVR has also been used to investigate the effect of healthy ageing on brain haemodynamics^7^.

However, the clinical adoption of hypercapnia based CVR mapping has so far been hindered by two main challenges: (i) relatively long scan times and (ii) long patient preparation times. Clinical MRI examinations are typically split into discrete diagnostic measurements of approximately 5 minutes in duration. However, currently CVR examinations are between 7 and 12 minutes in duration, meaning that compromises in diagnostic protocols must be made to accommodate such measurements. Preparation of patients adds an additional time penalty that is dependent on the means of gas administration and/or gas sampling. However, in this study we concentrate on the question of reducing the duration of the CVR examination.

Typically, hypercapnic gas challenges are presented in blocks interleaved with a baseline air breathing condition. One popular gas protocol used by researchers in Toronto involves the use of two hypercapnic blocks of differing duration; an initial short block followed by a second much longer block^8^. This protocol has also been used to derive information about the relative timing of the CVR response using Transfer Function Analysis (TFA)^9^. This timing information has been suggested to be reflective of regional variations in blood arrival time^10^ or in the rate at which the CVR response develops^9^. However, this block paradigm does not lend itself to easy scan time reduction as it typically relies on long hypercapnic blocks to minimise the effect of regional differences in vascular delays. Alternatively, in the limit of short blocks a sinusoidally varying stimulus has several useful properties^10^. Firstly, accurate synchronisation of the stimulus paradigm on a voxel by voxel basis is not required, as it is merely modelled as a phase shift of a sinusoid at the frequency of the stimulus. In this context the fitted amplitude of the sinusoid reflects CVR and the phase of the sinusoid provides similar timing information to TFA. With respect to the aim of reducing scan time, the sinusoidal stimulus protocol can be retrospectively reanalysed with fewer stimulus cycles, or fractions of cycles, since it is not the duration of the stimulus that is important but the stimulus frequency.

Therefore, the aim of this study was to develop a rapid CVR technique based on a sinusoidal hypercapnic stimulus. Comparison was made with a commonly applied block paradigm, which we term the Toronto protocol^8^. Both protocols produce estimates of vascular reactivity and the timing of this reactivity, which to prevent ambiguity we describe as CVR magnitude and CVR phase, respectively. Firstly, estimates of CVR magnitude and CVR phase were compared for the Toronto and Sinusoid protocols with the same scan duration. Secondly, the amount of data included in the analysis of the Sinusoid protocol was progressively reduced to investigate how rapidly CVR information can be acquired.

## Material and Methods

### Imaging

All imaging was performed on a Siemens Prisma 3T scanner (Siemens Healthineers, Erlangen, Germany) using the body transmit coil and the vendors 32-channel receive coil. This study was approved by the Medical Sciences Interdivisional Research Ethics Committee (MS IDREC) subcommittee of the Central University Research Ethics Committee (CUREC) at the University of Oxford (approval number: MSD-IDREC-C2-2014-041). Research was conducted in accordance with the Good Clinical Practice guidelines specified by the University of Oxford Clinical Trials and Research Governance unit. Ten healthy volunteers (age range 19 – 21, 5 female) were recruited and informed written consent obtained. Functional imaging consisted of BOLD-weighted EPI images with the following pulse sequence parameters: repetition time (TR) 2 s, echo time (TE) 30 ms, field of view (FOV) 220 mm × 220 mm, matrix 64 × 64, slices 24, slice thickness 5 mm, slice gap 0.5 mm, flip angle 80°, GRAPPA 2. Each functional scan had a duration of 7 mins, resulting in the acquisition of 210 imaging volumes. A field map was acquired using a 2D Fast Low Angle Shot (FLASH) method with the following parameters: TR 378 ms, TE1/TE2 4.92 ms / 7.38 ms, FOV of 220 mm × 220 mm, matrix 64 × 64, slices 24, slice thickness 4.5 mm, slice gap 0.45 mm, flip angle 45°. Finally, a high resolution T_1_-weighted structural image was acquired for each subject using a 3D Magnetisation Prepared Rapid Acquisition Gradient Echo (MPRAGE) pulse sequence^11^ with the following parameters: TR 1.9s, TE 3.74 ms, FOV 174 mm × 192 mm × 192 mm, matrix 116 × 128 × 128, flip angle 8°, inversion time (TI) 904 ms.

Details on how to access the imaging data that underlies this study can be found in Appendix A.

### Respiratory Stimulus

Hypercapnia challenges were delivered by a computer controlled gas blender (RespirAct™ Gen 3, Thornhill Research Inc., Toronto, Canada) that implements a prospective algorithm for the targeting of blood gases^12^. Subjects were fitted with a sequential gas delivery (SGD) breathing circuit, which was sealed to the face using adhesive tape (Tegaderm, 3M Healthcare, St. Paul, Minnesota, USA) to prevent the entrainment of room air. Subjects were then asked to sit comfortably outside the scanner whilst the prospective algorithm was calibrated. The subject’s specific resting end-tidal carbon dioxide and oxygen partial pressures (PetCO_2_ and PetO_2_, respectively) were first determined by averaging the first ten or so breaths into the breathing mask. The calibration consisted of initiating a baseline targeted at the subject’s specific resting PetCO_2_ and PetO_2_. Initial estimates of the subject’s VCO_2_ and VO_2_ (their resting production/consumption of CO_2_/O_2_) based on height, weight and age were refined until a constant baseline was achieved with minimal drift. A brief presentation of the sinusoidal hypercapnic stimulus was then administered to the subject to prepare them for the sensations they might encounter whilst in the scanner.

Gas stimulus protocols (Fig. 1a) were tailored to each subject’s resting PetCO_2_ and PetO_2_ baselines. Modulations in PetCO_2_ were targeted relative to baseline, whilst PETO_2_ was targeted to remain constant at baseline throughout. The Toronto gas protocol consisted of two blocks of hypercapnia; one of 45 s duration and the other 120 s. The former was preceded by a 60 s baseline and the latter by a 90 s baseline, with a final baseline period of 105 s completing the 7 minute protcol^13^. The sinusoidal gas protocol was composed of seven sinusoidal cycles, each with a period of 60 s. The amplitude was set to vary from baseline PetCO_2_ to 10 mmHg above baseline. In this implementation, the prospective algorithm is very tolerant of increases in ventilation rate compared with the baseline level during calibration, but not reductions in ventilation. Therefore, subjects were coached to maintain their ventilation rate over the scanner intercom should it be observed to have dropped below baseline levels.

**Figure 1.**
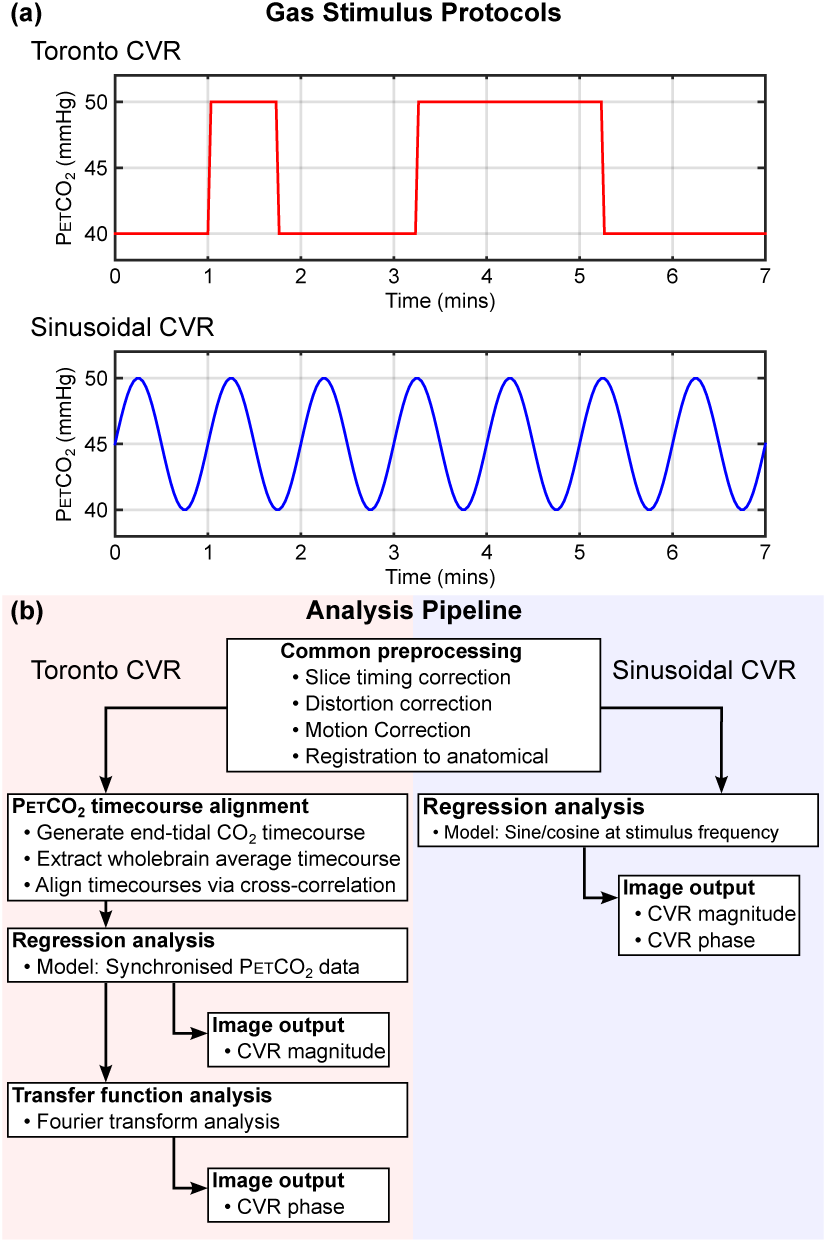
Data analysis pipeline for Toronto and Sinusoid protocols. It is important for the end-tidal CO_2_ regressor to be well-aligned with the BOLD data in the Toronto protocol to avoid underestimating changes in CVR Magnitude. The Sinusoid protocol analysis is inherently insensitive to such time delays and does not require this step.

During the presentation of each 7 minute stimulus protocol, imaging was performed using the 7 minute BOLD-weighted EPI pulse sequence described above. Synchronisation of the respiratory and imaging protocols was manually initiated.

### Image Analysis

The basic image analysis pipeline is summarised in Fig. 1b. Imaging data from the two stimulus protocols were first submitted to a common pre-processing stage including motion correction^14^, fieldmap based EPI unwarping^15^, slice timing correction, removal of non-brain tissue^16^ and spatial smoothing with a 5 mm FWHM kernel^17^. High pass temporal filtering was also performed with cut-off values of 100 s and 200 s for the Sinusoid and Toronto protocols, respectively. Images from the Toronto protocol were registered to MNI space via the subject’s structural image^14,18^. This registration was used to transform the MNI structural atlas^19^ and segmented structural images^20^ to functional space. Finally, the Sinusoid protocol images were registered to the Toronto protocol images.

The FMRIB Expert Analysis Tool (FEAT) was used to perform model-based multiple regression analysis^21^. Motion parameters were not included in this analysis. For the Sinusoid protocol, the model was defined by sine and cosine terms with time periods of 60 s (equivalent to 16.7 mHz). For the Toronto protocol, the model was defined by the measured PetCO_2_ change during the challenge experienced by each subject. This PetCO_2_ timecourse must first be synchronised with the associated BOLD signal changes in the brain, due to intersubject differences in blood arrival time. This was achieved by extracting the mean whole brain BOLD signal change, using the mask generated during the brain extraction process, and finding the maximum cross-correlation with the PetCO_2_ data interpolated to the BOLD sampling frequency. High pass temporal filtering with the same cut-off as the imaging data was then applied and the temporal derivative included as an additional regressor.

From this analysis several outputs can be produced. Statistical maps were generated and thresholded at the voxel level with a corrected voxel P threshold of 0.05. The effect of the sine and cosine terms in the sinusoidal analysis were combined by using an F-test, whereas for the Toronto analysis a single T-test was used. Maps of CVR magnitude were calculated from the parameter estimates (PE) of the GLM analysis using Eq. (1) for the Sinusoid protocol and Eq. (2) for the Toronto protocol, where S_mean_ is the mean BOLD signal baseline and ΔPetCO_2_ is the change in end-tidal partial pressure of CO_2_. Normalisation by ΔPetCO_2_ is included to control for difference in the stimulus magnitude across subjects.

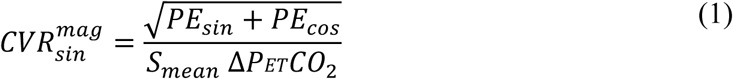

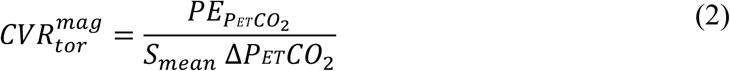

Despite the use of a GLM analysis in this work, in contrast to the Fourier analysis used previously^10^, it is still possible to calculate maps of CVR phase from the sinusoidal protocol by using Eq. (3).

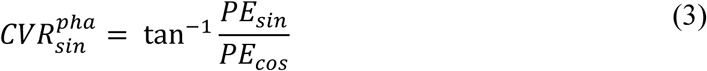

To provide CVR phase estimates that are comparable with the Toronto protocol, maps were produced using the whole brain phase as a reference i.e. equivalent to the synchronisation performed for the Toronto protocol. This was achieved by extracting a whole brain BOLD signal timecourse and estimating the CVR phase using Eq. (3). This global phase value was then subtracted from the CVR phase estimates from each voxel.

For the Toronto protocol Transfer Function Analysis (TFA)^9^ was used to estimate CVR phase. Since this is a Fourier based method, and as such no standard tools exist, it was implemented in MATLAB (Mathworks, Natick, MA, USA). The normalised transfer function H(f) is calculated by dividing the cross power spectral density (CPSD) of the BOLD and PetCO_2_ timecourses by the power spectral density (PSD) of the BOLD timecourse. Both the BOLD and PetCO_2_ timecourses were demeaned prior to spectral analysis.

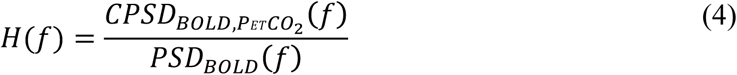

CVR phase can be calculated from the real and imaginary parts of H(f), denoted by subscript *r* and *i*, at the frequency of interest. This was chosen to be 10 mHz, consistent with previous work with the same stimulus protocol^9^.

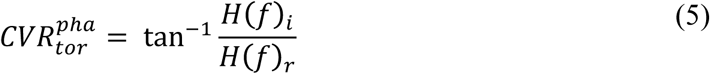

The use of a GLM analysis also enables the errors in measurements of CVR magnitude and CVR phase to be estimated. This can be achieved by propagating the errors in the GLM parameter estimates and residuals through to the final CVR estimate. The standard deviation in the CVR magnitude 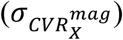 can be calculated using Eq.(6), where *PE_X_* and 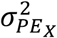 are the parameter estimate and the variance in that estimate for protocol X and 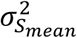 is the variance in the mean BOLD signal baseline.

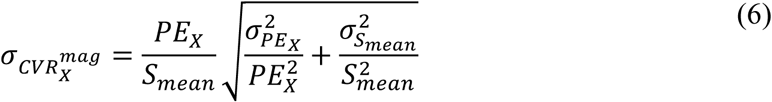

Since the Sinusoid protocol uses a pair of parameter estimates to calculate CVR magnitude the variance in those estimates must first be combined using the equation below before application of Eq.(6), where 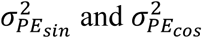 are the variance in the sine and cosine parameter estimates.

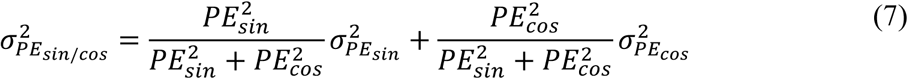

However, the absolute error in CVR magnitude does not take into account the large differences in the scale of CVR magnitude particularly between grey and white matter. Under these conditions it is preferable to represent the standard deviation as a fraction of CVR magnitude by calculating the relative standard deviation (RSD).

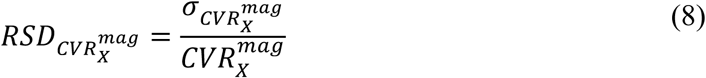

Estimating the variance in the CVR phase measurement was only possible for the Sinusoid protocol due to the use of TFA in the Toronto protocol.

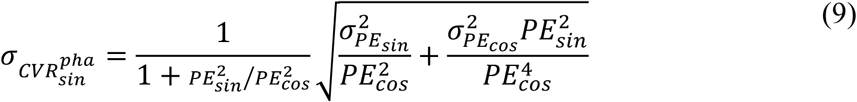

Details of how to access the code that was used to perform this analysis can be found in Appendix A.

### Stimulus performance

The normocapnic PetCO_2_ (PetCO_2,norm_) and the peak change in PetCO_2_ (∆PetCO_2_) were measured for each protocol and subject. First, the automated end-tidal picking routine was manually inspected and corrected where necessary, using software provided with the RespirAct, to ensure accurate PetCO_2_ estimation. Linear regression was then performed using a prototypical model of the stimulus (i.e. a boxcar temporally aligned with the PetCO_2_ data or sine/cosine pair at 16.7 mHz) and a constant term; the former representing ∆PetCO_2_ and the latter PetCO_2_,_norm_.

### Comparison between Toronto and Sinusoid protocols

Investigations were made to compare the sensitivity and equivalence of the two protocols. Firstly, the number of statistically significant voxels that passed the corrected voxel level threshold (p<0.05) were counted for each subject and protocol. In order to control for potential differences in subject motion between protocols the mean motion as calculated during the motion correction step was extracted^14^. Mean motion represents the mean-root-mean square displacement between adjacent time points and has been shown to be associated with reduced temporal signal to noise ratio (SNR) in fMRI experiments^22^. It is important that the effective temporal SNR is fairly constant across protocols in order to isolate the effect of different stimulus paradigm rather than difference in noise characteristics.

Next the equivalence of the CVR estimates produced by the Toronto and Sinusoid protocols was explored. Cortical ROIs were taken from the co-registered MNI atlas and further refined using a grey matter mask (grey matter segmentation partial volume estimate thresholded at 0.5). The mean and standard deviation of CVR magnitude and phase values extracted from these ROIs were calculated for each subject. Systematic differences between the estimates of CVR magnitude and phase from each of the protocols were assessed by fitting a simple model: linear slope and intercept.

### Effect of reducing Sinusoid protocol scan duration

The effect of reducing the amount of data used to measure the CVR parameters using the Sinusoid protocol was explored. This was achieved by truncating the data set at scan durations between 1 minute (30 volumes) and 6 minutes (150 volumes) in steps of 1 minute (30 volumes). The full analysis, including pre-processing and GLM analysis, was performed for each truncated dataset, and maps of CVR magnitude and phase were generated. High pass temporal filtering at 100 s was disabled for the 1 minute scan duration.

The impact of scan time reduction was investigated at the level of whole brain grey matter. An ROI was generated based on voxels in the Toronto protocol that met the statistical threshold and was then refined using a grey matter mask (grey matter segmentation partial volume estimate thresholded at 0.5). The mean and standard deviation of the CVR magnitude and phase values in this ROI were calculated for each scan duration and subject. In addition, the number of statistically significant voxels from the GLM analysis were calculated for each scan duration and subject. These results were subjected to a two-way ANOVA test to investigate the null hypothesis that the group mean values were equal. In the case that the group mean values were found to be significantly different, pairwise comparison was performed using the Tukey-Kramer method (honest significance difference test) to consider which pairs of group means were significantly different. The Tukey-Kramer method controls for multiple comparisons by correcting for the family wise error rate. In this sense it is less conservative than multiple T-tests with Bonferroni correction.

## Results

Both stimulus protocols yielded reliable and accurate respiratory challenges. Table 1 details this performance by listing the baseline PetCO_2_ and stimulus induced ∆PetCO_2_. Although within the accuracy of the end-tidal gas targeting system, on average baseline PetCO_2_ was 0.9 mmHg higher and ∆PetCO_2_ was 0.7 mmHg lower for the Toronto protocol compared with the Sinusoid protocol. Paired two tailed T-tests showed that these differences were significant at p<0.001 and p<0.01, respectively. Although the target of 10 mmHg PetCO_2_ change was not attained, all subjects experienced similar PetCO_2_ changes within a standard deviation of 1.2 mmHg.

**Table 1.** Respiratory stimulus performance across protocols as defined by baseline normocapnic PetCO_2_ (PetCO_2,norm_) and the change in PetCO_2_ due to the stimulus
(ΔPetCO_2_).

| Subject ID | Toronto protocol |  | Sinusoid protocol |  |
| --- | --- | --- | --- | --- |
| | $\text{PETCO}_{2,\text{norm}}$ | $\Delta\text{PETCO}_2$ | $\text{PETCO}_{2,\text{norm}}$ | $\Delta\text{PETCO}_2$ |
| 01 | 38.6 | 7.0 | 37.8 | 7.9 |
| 02 | 37.9 | 7.1 | 37.1 | 8.0 |
| 03 | 43.1 | 6.2 | 41.9 | 6.9 |
| 04 | 43.8 | 5.0 | 42.2 | 5.9 |
| 05 | 40.4 | 7.4 | 39.5 | 7.5 |
| 06 | 31.0 | 6.0 | 29.9 | 6.6 |
| 07 | 41.2 | 5.3 | 40.5 | 6.2 |
| 08 | 40.6 | 7.9 | 40.0 | 8.0 |
| 09 | 38.7 | 8.0 | 38.5 | 8.5 |
| 10 | 38.7 | 7.7 | 37.4 | 9.7 |
| Mean | 39.4 | 6.8 | 38.5 | 7.5 |
| SD | 3.5 | 1.1 | 3.5 | 1.2 |

In order to compare the sensitivity of the two protocols the number of voxels that passed the voxel level statistical threshold were counted. Table 2 displays these results for the full 7 minute data sets, as well as two examples of the truncated Sinusoid protocol data. The mean motion is also tabulated here to demonstrate the impact of motion on the statistical maps. Mean motion is fairly consistent between protocols and truncated data sets, with one exception. Subject 10 shows considerably higher mean motion for the full Sinusoid protocol compared with the Toronto protocol or truncated data sets. Further investigation revealed large translations in the last two stimulus cycles. Since this would lead to a varying error contribution in the GLM analysis across truncated data sets, this subject was excluded from further analysis. For the remaining subjects, a paired two-tailed T-test showed that the number of statistically significant voxels was insignificantly different between the Toronto and Sinusoid protocols (p=0.36).

**Table 2.** Number of voxels above the voxel level statistical threshold for the 7 minute datasets of each protocol and the Sinusoid protocol truncated at 5 minutes and 3 minutes. The mean motion, defined as average root mean square displacement between adjacent time points, for each data set is presented alongside voxel count to highlight the effect of subject motion.

| Subject<br>ID | Toronto protocol |  | Sinusoid protocol |  |  |  |  |  |
| --- | --- | --- | --- | --- | --- | --- | --- | --- |
|  | 7 mins |  | 7 mins |  | 5 mins |  | 3 mins |  |
|  | #<br>voxels | Motion | #<br>voxels | Motion | #<br>voxels | Motion | #<br>voxels | Motion |
| 01 | 18995 | 0.11 | 17089 | 0.10 | 14735 | 0.10 | 10345 | 0.11 |
| 02 | 15650 | 0.13 | 17054 | 0.10 | 15969 | 0.10 | 15380 | 0.11 |
| 03 | 17820 | 0.14 | 15104 | 0.15 | 14221 | 0.14 | 13695 | 0.13 |
| 04 | 9590 | 0.20 | 10174 | 0.20 | 8736 | 0.20 | 5013 | 0.19 |
| 05 | 16828 | 0.09 | 17185 | 0.09 | 15764 | 0.09 | 13941 | 0.09 |
| 06 | 5090 | 0.27 | 7106 | 0.29 | 6895 | 0.29 | 4785 | 0.22 |
| 07 | 15507 | 0.20 | 14643 | 0.24 | 12490 | 0.22 | 12081 | 0.21 |
| 08 | 19356 | 0.23 | 12400 | 0.30 | 10986 | 0.32 | 8070 | 0.34 |
| 09 | 19333 | 0.08 | 19507 | 0.12 | 17852 | 0.12 | 15109 | 0.10 |
| 10 | 19630 | 0.19 | 6767 | 0.40 | 15776 | 0.21 | 12890 | 0.22 |
| Mean | 15780 | 0.16 | 13703 | 0.20 | 13343 | 0.18 | 11131 | 0.17 |
| SD | 4814 | 0.06 | 4437 | 0.11 | 3509 | 0.08 | 3945 | 0.08 |

Examples of the main outputs of the analysis are displayed for both protocols in Fig. 2, consisting of unthreholded Z-statistic maps, CVR magnitude, CVR magnitude RSD, CVR phase and CVR phase standard deviation (SD). Both protocols can be seen to produce qualitatively similar results. A reduced Z-statistic value is observed in the Sinusoid protocol compared with the Toronto protocol (Fig. 2a/e). However, similar grey matter to white matter contrast is observed across CVR magnitude (Fig. 2b/f) and phase (Fig. 2d/h) maps. Similarly estimates of the error in these parameters demonstrate increased variance in white matter compared with grey matter. To investigate the quantitative equivalence of the protocols Fig. 3 plots cortical ROI estimates of CVR magnitude (Fig. 3a) and CVR phase (Fig. 3b) from both methods. Both CVR parameters are significantly correlated across protocols; CVR magnitude p<0.001 and CVR phase p<0.05. Linear regression across the nine remaining subjects produced a slope of 0.97±0.05 for CVR magnitude and 0.34±0.15 for CVR phase. Per subject estimates of the slope reveal that the group mean is not significantly different from unity for CVR magnitude (p=0.98, Fig. 3c) or for CVR phase (p=0.13).

**Figure 2.**
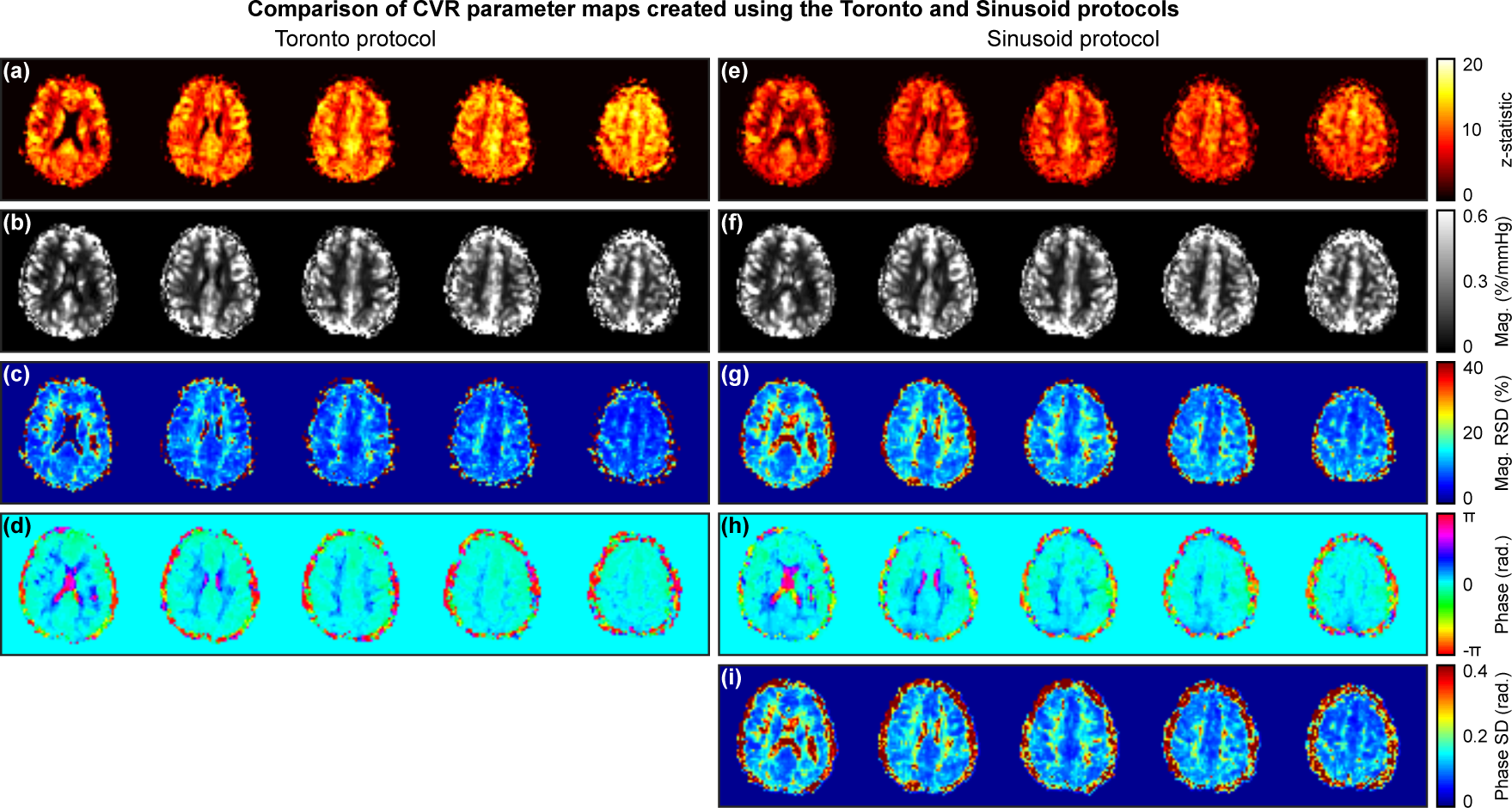
Example CVR maps from a single subject (Subject 01). A subset of five contiguous slices are displayed superior to the lateral ventricles. Maps (a)-(d) were produced using the Toronto protocol and (e)-(i) using the Sinusoid protocol. Maps were generated to display (a/e) statistical correlation with the CO_2_ stimulus (Z-statistic), (b/f) CVR magnitude (normalised by the change in PetCO_2_), (c/g) relative standard deviation (RSD) of the CVR magnitude estimate, (d/h) CVR phase and (i) standard deviation of the CVR phase estimate. Note: It is not possible to generate maps of the error in CVR phase for the Toronto protocol.

**Figure 3.**
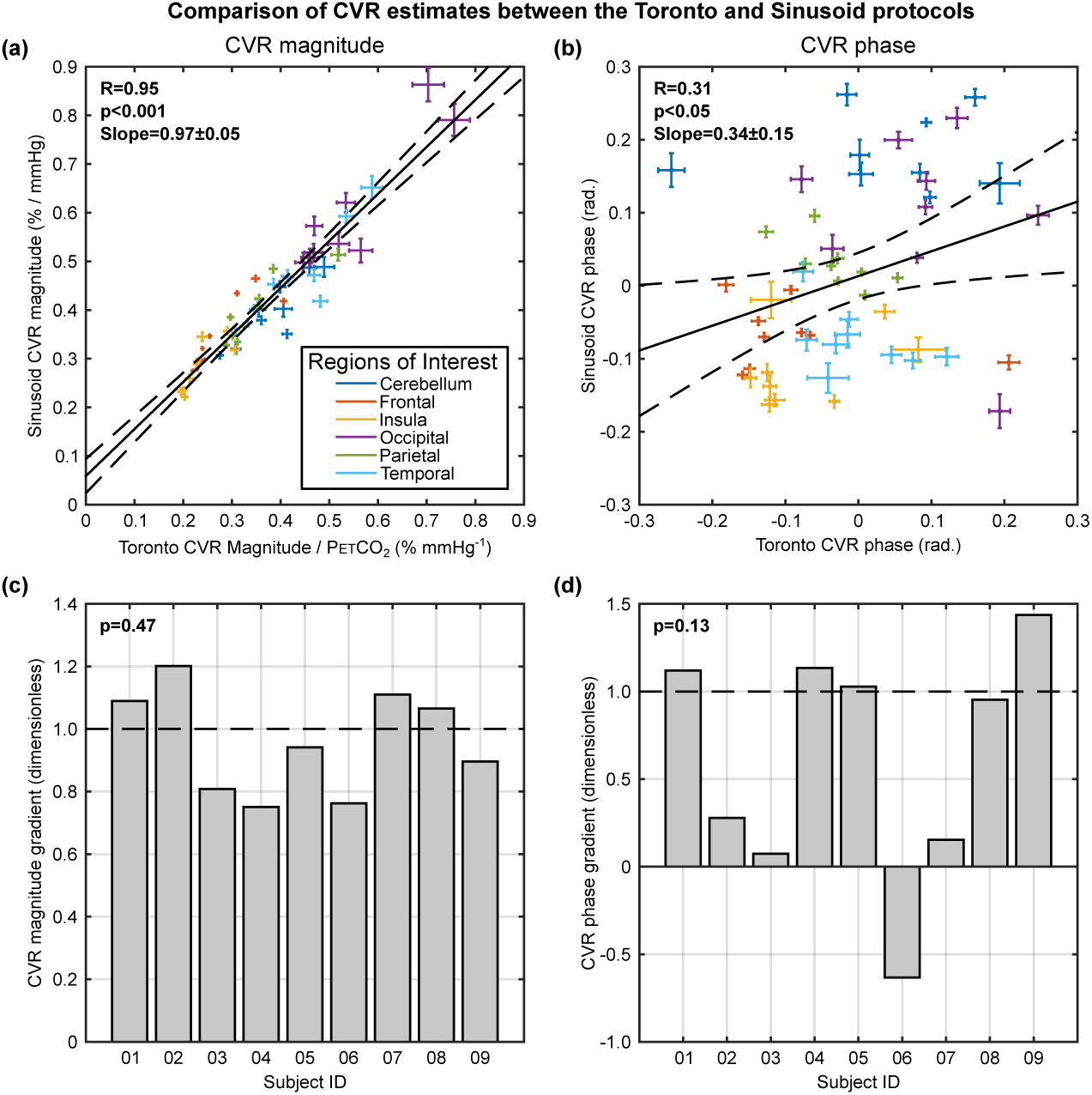
Estimates of (a) CVR magnitude and (b) CVR phase were compared on the basis of cortical regions of interest at the level of the cerebral lobes. Both measurements were found to be significantly correlated across protocols. To assess the equivalence of the two protocols a simple linear model was fitted for each subject. The mean and standard deviation of the slope was found to be 0.96±0.17 and 0.62±0.68 for (c) the CVR magnitude and (d) the CVR phase, respectively. Neither were found to be significantly different to unity, indicating that estimates from the protocols are equivalent.

An example of the effect of reducing scan duration is displayed in Fig. 4. Two examples of the truncated Sinusoid protocol (5 minutes and 3 minutes) are presented alongside the full 7 minute acquisition. Values of the Z-statistic can be seen to reduce with the amount of data included in the analysis (Fig. 4a-c). CVR magnitude maps look qualitatively very similar across scan durations (Fig. 4d-f), whilst CVR phase maps show some small differences when only 3 minutes of data are used (Fig. 4g-i). This is further explored across all subjects in Fig. 5. The group mean of the mean CVR magnitude across the ROI (Fig. 5a) was seen to increase slightly with reduced scan time. A two-way ANOVA test showed that not all group means were equal (p<0.001), but a paired multiple comparisons analysis showed that only the 1 minute duration was significantly different (p<0.001) to the full data set. A similar increase was observed for the group mean standard deviation of the CVR magnitude (Fig. 5b). Again group means were not all equal (p<0.001) and only the 1 minute duration was significantly different (p<0.001) to the full data set. The group mean of the mean CVR phase displayed little variation (Fig. 5c) and was not significant (p=0.99). The group mean of the standard deviation of the CVR phase showed the largest increase with time reduction (Fig. 5d). The group means were significantly different (p<0.001) and the 1 minute (p<0.001) and 2 minute (p<0.01) scan durations were significantly different to the full data set. The group mean number of statistically significant voxels (Fig. 5e) reduced with scan time and were significantly different (p<0.001). Pairwise comparison revealed significant differences between the 1, 2 and 3 minute durations (p<0.001) and the full data set. Finally, for comparison Fig. 5f scales the group means of the standard deviation of the CVR magnitude and phase to the full data set, demonstrating a larger effect of scan time reduction on CVR phase than CVR magnitude.

**Figure 4.**
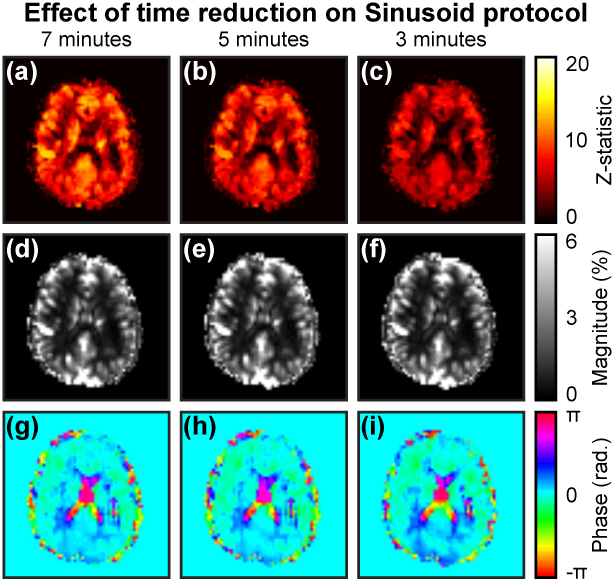
Example single slice of CVR maps from the Sinusoid protocol as scan duration is reduced (Subject 01) for (a)-(c) statistical correlation, (d)-(f) CVR magnitude and (g)-(i) CVR phase.

**Figure 5.**
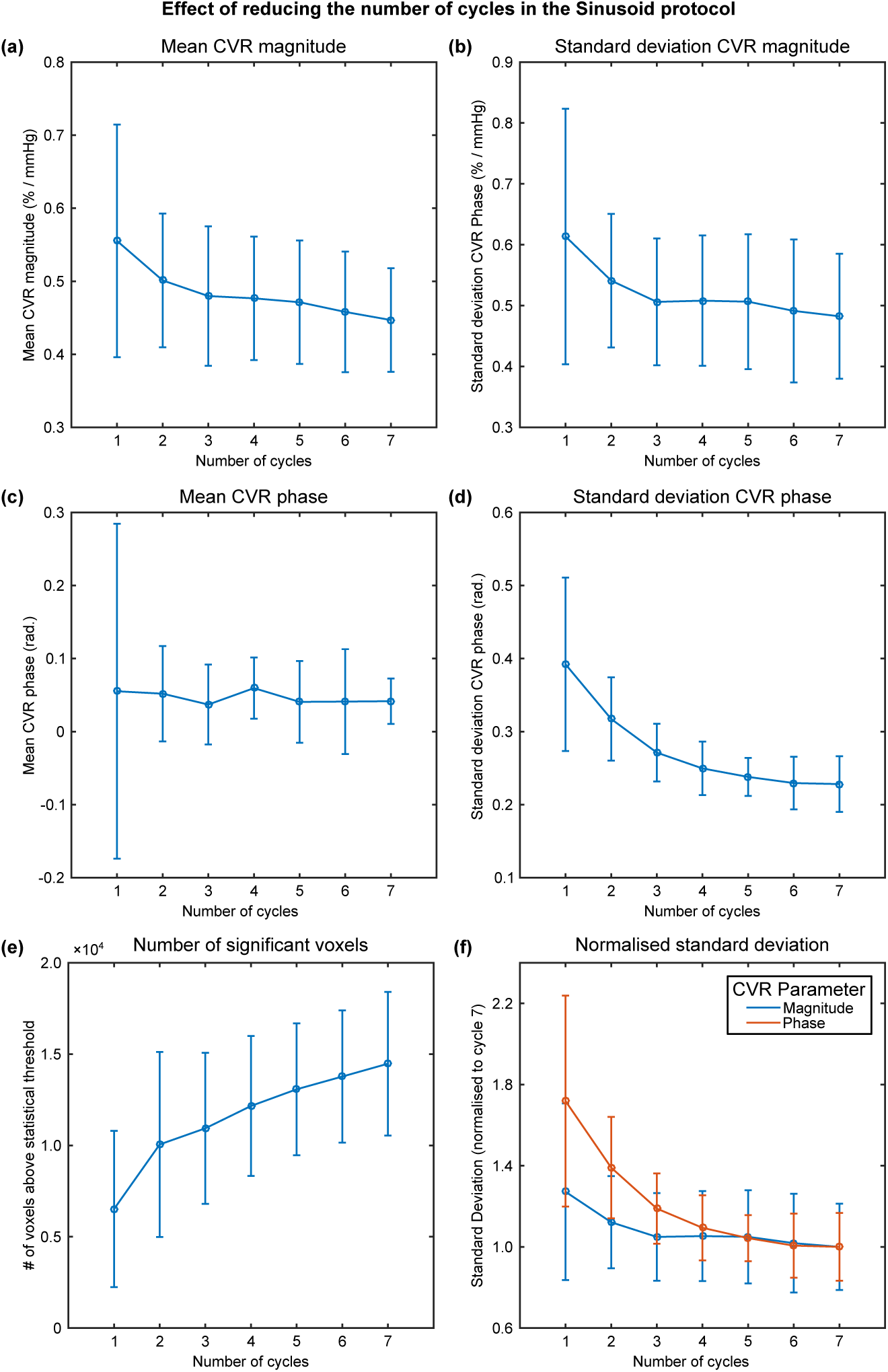
Analysis of the effect of reducing the scan duration of the Sinusoid protocol for (a/b) CVR magnitude, (c/d) CVR phase and (e) the number of voxels above the voxel level statistical threshold. Analysis was performed for a grey matter region of interest (ROI) where the mean and standard deviation of the estimates within the ROI were calculated. The standard deviation of the CVR magnitude and phase are compared directly (f) by normalising to the 7 minute scan duration.

## Discussion

CVR mapping using MRI is a promising clinical tool for stress testing cerebral blood vessels. However, there are several aspects of current CVR mapping techniques that limit their clinical application. In this study we concentrated on minimising the scan duration in order to improve the compatibility of this technique with clinical workflows. By utilising a sinusoidally modulated hypercapnia challenge we were able to retrospectively examine the effect of reducing the scan duration. In turn we were able to demonstrate that good quality CVR maps can be acquired in around 3 minutes, rather than the more typical 7 minutes. Furthermore, we demonstrated that CVR maps generated using this sinusoidal stimulus are equivalent to the more conventional block paradigm.

### Stimulus performance

Gas stimulus protocols were tailored to each subject’s individual baseline PetCO_2_. This baseline was maintained within 1 mmHg between the protocols, but was statistically significantly different none the less. Since the vascular response to carbon dioxide is sigmoidal in form^23^ it is possible that the two protocols were performed on different parts of the sigmoid curve. However, in practice maintaining baseline PetCO_2_ to a higher accuracy than 1 mmHg is challenging. Both stimulus protocols elicited PetCO_2_ changes of the order of 7 mmHg, which should place the vascular response within the linear regime of the sigmoidal vascular response^23^. A statistically significant difference in ∆PetCO_2_ was observed, although well within the accuracy of the gas delivery system at 0.7 mmHg. Unfortunately, subject tolerance of both stimulus protocols was not recorded. However, none of the subjects reported discomfort from either stimulus protocol.

### Comparison between Toronto and Sinusoid protocols

Before discussing the results of the protocol comparison it is useful to consider the advantages and disadvantages of the two stimulus protocols. As a more conventional block paradigm, the Toronto protocol is an easier stimulus to implement. However, synchronisation of the measured PetCO_2_ experienced by the subject with the BOLD data is critical to the accurate estimation of magnitude of the CVR change. Whilst this is practical when synchronising with the whole brain average BOLD signal, the presence of local variations in vascular delays opens up the possibility of the underestimation of CVR magnitude, or the measurement of spurious negative signal changes^9^. This can be mitigated by the use of long blocks of hypercapnia, but therefore limits how short the total acquisition can be. In contrast sinusoidal stimuli have seen only limited application^24^. To be clinically practical, computer controlled gas delivery is desirable, although a manual implementation has also been demonstrated^25^. In the previous implementation a Fourier analysis approach was used based on in-house analysis software^10^. However, in this study we reframe the analysis in order to use free and widely available fMRI analysis tools^15^. One of the inherent advantages of the Sinusoid protocol is its insensitivity to local vascular delays. This is due to the fact that any sinusoid with arbitrary amplitude and phase can be represented by the sum of a sine and a cosine with different amplitudes. In this context only the frequency of the stimulus is important, rather than the timing. This enables the short time-period stimuli used in this work, which would otherwise result in underestimates, or artefactual negative values, of CVR magnitude. Finally, simple sine and cosine functions were used as regressors in the GLM analysis, rather than the measured PetCO_2_ timecourse. Despite this, the number of statistically significant voxels (Table 2) was not significantly different to the Toronto protocol across the group. This approach simplifies the analysis as the PetCO_2_ data does not need to be transferred to the analysis computer and synchronised with the BOLD data.

Maps of CVR magnitude were found to have qualitatively similar features (Fig. 2b/f). Furthermore, quantitative analysis comparing regional estimates of CVR magnitude at the level of the cerebral lobes demonstrated significant correlation between protocols (Fig. 3a) and equivalence of the absolute values (Fig. 3c). By utilising estimates of the variance in the parameter estimates and the residuals from the GLM analysis it was possible to evaluate the standard deviation of the CVR magnitude measurement. By presenting this as the relative standard deviation (RSD), with respect to the mean, the large degree of heterogeneity in CVR magnitude is controlled, revealing more noisy estimates in white matter (Fig. 2c/g). However, the standard deviation in CVR magnitude, as estimated using Eq. (6), should be viewed with caution as it relies on the variance in S_mean_ from the residuals. The residuals include contributions from motion and physiological noise that aren’t included in the GLM model. Therefore, large amounts of motion will result in larger values of RSD, in effect being categorised as noise.

Maps of CVR phase also showed clear similarities between methods (Fig. 2d/h). Quantitative analysis showed that the protocols were significantly correlated (Fig. 3b). However, the slope between the protocols varied in the range −0.63 to 1.44 and was trending towards being significantly different to unity (the value expected for equivalence). This disparity may be due to the properties of phase, particularly the incorrect unwrapping of phase values close to ±2π. It is also possible that CVR phase behaves differently between the protocols given the large difference in stimulus duration between the Toronto (45–120 s above baseline PetCO_2_) and the Sinusoid protocol (30 s above baseline PetCO_2_ in each 60 s cycle). Maps of the standard deviation of the CVR phase measurement were estimated for the Sinusoid protocol by propagating the errors in the parameter estimates (Fig. 2i). RSD was not used because of the relative nature of phase i.e. the reference phase is arbitrary. It is not possible to estimate the standard deviation of the Toronto CVR phase due to the use of TFA, although an analysis of the effects of noise was included in the original paper^9^.

### Effect of reducing Sinusoid protocol scan duration

Rather than acquire multiple scans with different durations, the same 7 minute protocol was truncated to investigate the effect of reducing scan duration. From a qualitative perspective, whilst the values of the Z-statistic are reduced with decreasing scan duration, there appears to be relatively little effect on maps of CVR magnitude at scan durations of 3 or 5 minutes, and only small differences in the 3 minute CVR phase map (Fig. 4). Quantitative analysis confirmed that the number of above threshold voxels in the Z-statistic maps was significantly different in the 1, 2 and 3 minute scan durations when compared with the full data set. Furthermore, the results in Fig. 5e are consistent with the expected reduction in sensitivity demonstrated for fMRI analysis^26^. CVR magnitude was only minimally affected, consistent with the qualitative assessment above, with only the 1 minute scan duration found to be significantly different to the full data set for both the mean and standard deviation across the ROI. Estimates of CVR phase appear to be more strongly affected by scan time reduction. Whilst the mean over the ROI was not significantly different as a function of scan duration, the standard deviation was significantly different for the 1 and 2 minute durations. This increased sensitivity of CVR phase to scan time reduction, relative to CVR magnitude, was highlighted by normalising Fig. 5b and Fig. 5d to their 7 minute values. The standard deviation of CVR phase over the ROI clearly increases much more rapidly for CVR phase than CVR magnitude, necessitating longer scan duration where CVR phase is a priority. In summary, the results would suggest that scan time could be reduced to be in the range of 3 to 5 minutes with minimal effect on the resulting maps of CVR.

### Limitations and future work

In the current implementation of this technique, long patient preparation times remain the largest barrier to clinical adoption. Whilst it was not the aim of this study to tackle this problem, it does deserve further consideration. To maintain a constant PetCO_2_ baseline with minimal drift using the prospective end-tidal gas targeting system used in this study, the system must be calibrated to each individual. In our hands this process takes about 30 minutes, including application of the mask, although it may be shorter for more experienced users. However, there is a great deal of scope for time reduction by automating this calibration step, which will likely be implemented in future hardware developments. In the short term, using the current hardware, it should still be possible to realise shorter patient preparation times, albeit at the expense of a small amount of PetCO_2_ baseline drift. As noted above, the gas targeting system provides an initial setup based on the height, weight and age of the subject. The calibration process enables this setup to be refined. However, the initial estimate results in only small amounts of drift in many cases. Small amounts of drift are removed by the high-pass temporal filter and will not affect estimates of CVR so long as the vasculature remains in the linear regime of the sigmoidal vascular response. Further work is required to investigate this approach.

## Conclusions

In this study we have demonstrated that a sinusoidal hypercapnia stimulus provides equivalent information about CVR to a conventional block paradigm. Furthermore, we have shown that the scan duration of the Sinusoid protocol can be drastically reduced with only a small effect on the resulting CVR maps. This development should enable scan durations between 3 and 5 minutes and has the potential to enable more widespread clinical adoption.

## Appendix A. Supplementary data

The raw data that underpins this work can be accessed via the Oxford Research Archive repository, doi: https://doi.org/10.5287/bodleian:Xk48adQAO. Furthermore, the code to perform the analyses on these data can be accessed via the Zenodo repository, doi: http://doi.org/10.5281/zenodo.203128.

## Funding

This work was supported by the Engineering and Physical Sciences Research Council [grant number EP/K025716/1]. DPB also received salary support from Cancer Research UK.

## Declaration of conflicting interests

The author(s) declared no potential conflicts of interest with respect to the research, authorship, and/or publication of this article.

## Author’s contributions

NPB and DPB designed the study. NPB, JWH and DPB performed the research. NPB analysed the data. NPB, JWH and DPB wrote the paper.

